# Mapping the common and distinct neural correlates of visual, rule and motor conflict

**DOI:** 10.1101/2020.11.19.388231

**Authors:** Bryony Goulding Mew, Darije Custovic, Eyal Soreq, Romy Lorenz, Ines Violante, Stefano Sandrone, Adam Hampshire

**Affiliations:** The Computational, Cognitive and Clinical Neuroimaging Laboratory (C3NL), Department of Brain Sciences, Imperial College London, United Kingdom; MRC Cognition & Brain Sciences Unit, University of Cambridge, United Kingdom; School of Psychology, University of Surrey, Guildford, United Kingdom

## Abstract

Flexible behaviour requires cognitive-control mechanisms to efficiently resolve conflict between competing information and alternative actions. Whether a global neural resource mediates all forms of conflict or this is achieved within domain-specific systems remains debated. We use a novel fMRI paradigm to orthogonally manipulate rule, response and stimulus-based conflict within a full-factorial design. Whole-brain voxelwise analyses show that activation patterns associated with these conflict types are distinct but partially overlapping within Multiple Demand Cortex (MDC), the brain regions that are most commonly active during cognitive tasks. Region of interest analysis shows that most MDC sub-regions are activated for all conflict types, but to significantly varying levels. We propose that conflict resolution is an emergent property of distributed brain networks, the functional-anatomical components of which place on a continuous, not categorical, scale from domain-specialised to domain general. MDC brain regions place towards one end of that scale but display considerable functional heterogeneity.

## Introduction

The human brain maintains stable, yet flexible, behaviour through the employment of cognitive control mechanisms (Rougier et al., 2005; Stokes et al., 2017). These mechanisms allow information processing to be adjusted relative to situational requirements (Dosenbach et al., 2008; Braver et al., 2009). Competing sources of information and alternative actions can have conflicting neural representations, which compromises task performance (Botvinick et al., 2001). Consequently, a prominent function of cognitive control is to efficiently resolve conflict, thereby enabling optimal behaviour.

The mechanisms underlying conflict resolution have been widely debated. Broadly speaking, there are two distinct classes of model. One class proposes that conflict resolution is domain-specific, with different brain regions specialised to process distinct sources of conflict (Egner et al., 2007; Notebaert and Verguts, 2008; Akçay and Hazeltine, 2011; Kim et al., 2012). From this perspective, multiple, independent control mechanisms are thought to operate in a conflict-driven manner, whereby the resolution of conflict from one source is independent of conflict from another (Egner, 2008; Kiesel et al., 2006). Evidence to support this domain-specific perspective has come from studies looking at behavioural conflict adaptation, where task performance on a given trial improves if the previous trial was ‘high conflict’ (Verbruggen et al., 2005; Notebaert and Verguts, 2008; Akçay and Hazeltine, 2011; Kim et al., 2012). It has been proposed that control mechanisms are upregulated upon initial exposure to conflict in a domain-specific manner, and there is a subsequent improvement in task performance (Botvinick et al., 2001).

Manipulating the conditions under which conflict adaptation occurs has been used to characterise the nature of these underlying mechanisms. For example, Akçay, and Hazeltine (2011) reported that conflict adaptation was evident when high conflict on the previous trial was within-type but not across-type. This indicates that conflictresolution mechanisms have a degree of dimension specificity. Neuroimaging evidence have also supported this hypothesis, showing activity in the superior parietal cortex for stimulus-based conflict vs. the ventral premotor cortex for response-based conflict (Egner et al., 2007; Kim et al., 2010).

In direct contrast to the localist perspectives is the hypothesis that a general cognitive system processes diverse sources of conflict (Cole and Schneider, 2007; Crittenden et al., 2016; Egner et al., 2007; Freitas et al., 2007; Kim et al., 2012). This hypothesis accords well with the broader globalist school of thought, which states that cognitive functions are rarely attributed to isolated brain regions. Instead, a non-discriminatory global cognitive control mechanism could flexibly adapt to resolve diverse sources of conflict (Cole and Scheider, 2007; Hsu et al., 2017; Kan et al., 2013). Brain regions commonly attributed a global role in conflict resolution include the dorsolateral prefrontal cortex (DLPFC), inferior frontal junction, cingulo-opercular network, dorsal premotor cortex, pre-supplementary motor area and adjacent dorsal anterior cingulate and intraparietal sulcus (Botvinick et al., 2001; Ambrosini and Vallesi, 2017; Li et al., 2017; Wu et al., 2020). These regions are consistent with ‘multiple-demand cortex’ (MDC) (Duncan 2001, Duncan, 2010), the set of brain regions that is most commonly activated across diverse cognitive tasks, that is, regardless of information type (Assem et al., 2020; Duncan, 2001; Duncan, 2010). This cortical volume is considered to support a ‘task set’ which is maintained throughout the task and is responsible for the coordination of processing strategies relative to demands (Melcher et al., 2008).

Although conflict adaptation studies often support a domain-specific perspective, some have reported cross-domain conflict improvements in task performance, which could indicate global control (Kan et al., 2013; Kunde and Wühr., 2006). For example, Kan and colleagues (2013) observed conflict adaptation effects between tasks that were within-domain (verbal to verbal) and those that were across-domain (perceptual to verbal), suggesting that conflict resolution can generalise across different conflict types. However, the domain-general hypothesis is primarily supported by neuroimaging data. Fan and colleagues (2003) analysed three tasks, each with a distinct source of conflict. The results indicated that a global network of regions contribute to conflict resolution of any type, but there was also some activity unique to each task. Similarly, Hsu et al., (2017) identified common activity patterns in response to conflict from multiple origins, suggesting influence from a domain-general system.

A limitation though is that conflict resolution mechanisms are assessed using a diverse tasks throughout the literature, which impedes comparisons between the two classes of model. Stimulus switching, task switching and factorial designs are common approaches (Egner, 2008). Thus, although the literature offers evidence for both local and global models of conflict resolution, there is no consensus on which is correct. Indeed, the two proposed classes of model are not necessarily mutually exclusive. It could be the case that MDC sub-regions are recruited for all types of conflict, but they operate in conjunction with local conflict resolution mechanisms, such as lateral inhibition (Erika-Florence et al., 2014; Hampshire & Sharp, 2015). Furthermore, if MDC subregions have greater proximity to different types of information input (Shashidhara et al., 2019), then this may be reflected in partial dissociations, that is, being recruited *en masse,* but to varying degrees dependent on the functional-anatomical locus of conflict.

Here, we adjudicate between these possibilities by using a novel functional magnetic resonance imaging (fMRI) paradigm designed to orthogonally manipulate conflict at different stages of the stimulus-rule-response process. The mixed block/event-related design used motion-coherence, relational rule and Go/No-Go manipulations to vary conflict at different stages of the stimulus-response process (**Figure 1**). The participants (14 female and 7 male) received no feedback during the task; therefore, the mapping rules were established by a process of instruction-based learning (Cole et al., 2013a; Hampshire et al., 2016 & 2019; Ruge and Wolfenstellar, 2013). By applying a fully factorial design with discrete switching and trial stages, we map the brain regions that are activated by increased conflict within specific domains whilst controlling for general difficulty and other confounding task demands. Contrasts and conjunction analyses are used to map distinct and common correlates of conflict resolution. Focused ROI analyses then test at a finer grain whether MDC sub-regions have uniform or dissociable sensitivities to conflict demands and determine if such functional dissociations are absolute or a matter of degree.

**Figure 1.**
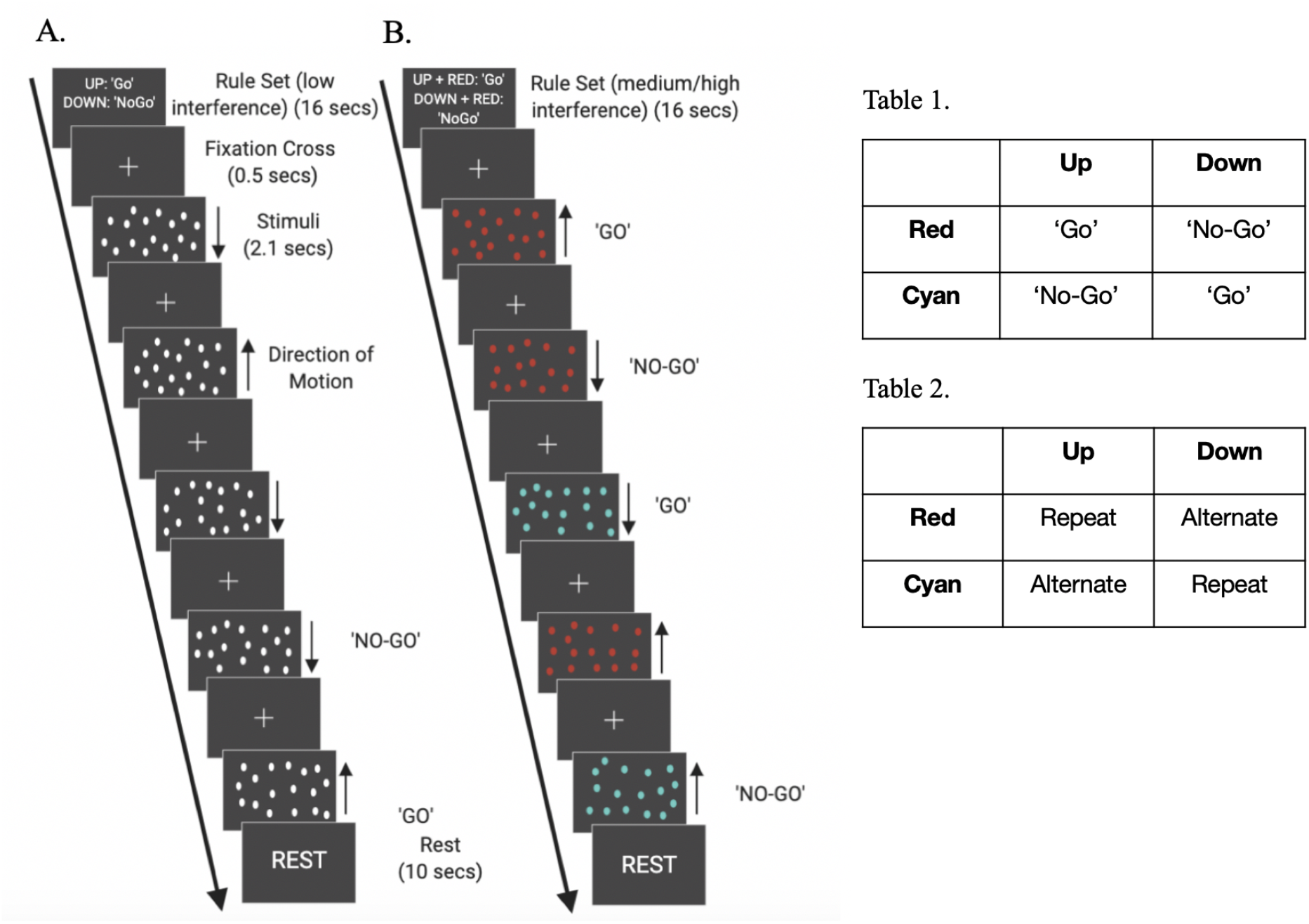
Examples of blocks within the task. Block 1 (A.) shows stimulus-response mapping rule level 1 (R1) and block 2 (B.) shows level 2/3 (R2/R3). Participants were initially shown an instruction slide for 16 seconds before being presented with ten stimuli, each with a duration of 2.1 seconds, separated by an inter-stimulus interval (fixation cross) of 0.5 seconds. The stimulus conditions at each trial required the participants to make a choice between response A or B (Go or No-Go). Each block finished with a 10 second rest. (A.) Instruction slide and example trials for blocks featuring R1 where colour is not a relevant stimulus feature. (B.) Instruction slide (incomplete) and example trials for blocks featuring R2/3 where both direction of coherent motion and colour are relevant. R3 includes an additional temporal order dependency. **(Table 1.)** Possible stimuli defined by both motion direction and colour, and the corresponding required responses for the ‘medium’ level along the rule dimension of the task space. **(Table 2.)**Possible stimuli including an additional temporal factor and the corresponding required responses for the ‘high’ depth of rule integration along the rule dimension of the task space.

## Results

### Performance accuracy decreases as stimulus and rule interference increase

Mean task performance during imaging acquisition was well above chance. Factorial ANOVAs with repeated measures were estimated to examine the main effects of stimulus level, rule level and response level and the interaction effect between factors on accuracy and response time. For response accuracy (RA), there was a significant main effect of stimulus level (F(1,20) =126.47; p<0.001; *d*= 1.184 stimulus 1>stimulus 2) and rule level (F(2,19) =32.43; p<0.001; *d*= 0.055 (rule 1>rule 2); *d*= 2.98 (rule 2>rule 3); *d*= 2.50 (rule 1>rule 3), but not for response level (F(1,20)=0.636; p=0.435; *d*=0.08 (response 1>response 2), as depicted in **Figure 2A**. There was a significant interaction between rule and response (F(2,19) =9.884; p=0.001), but not between stimulus and rule (F(2,19)=3.254; p=0.061) or stimulus and response (F(1,20)=1.313; p=0.265). The three-way interaction was non-significant (F(2,19)=2.066; p=0.154).

**Figure 2.**
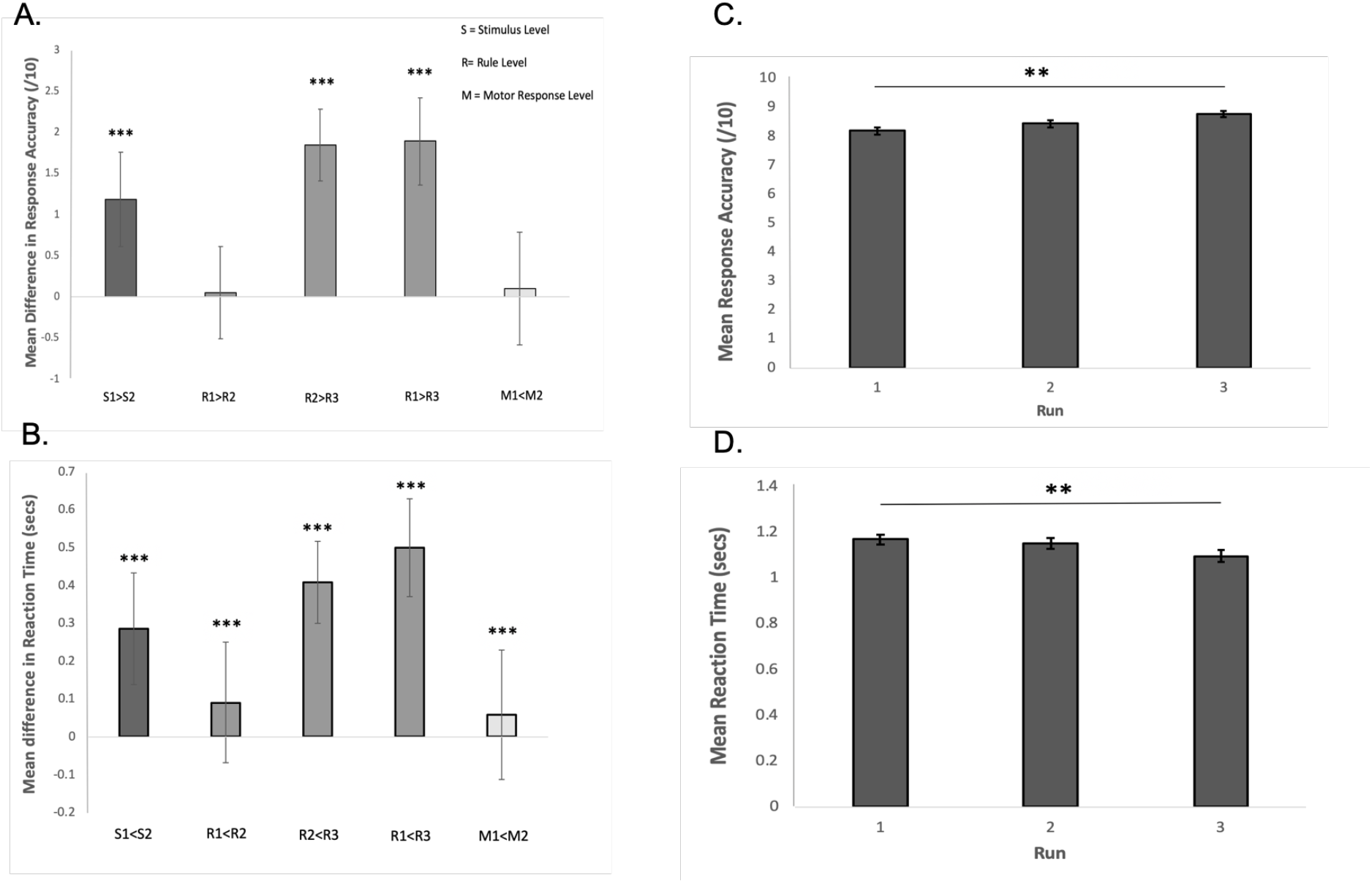
Behavioural results. (A) Difference in response accuracy between pairs of levels within the factorial design (Y scale is mean difference in accuracy count per block). During stimulus and rule interference, response accuracy is greater at a lower task difficulty. (B) Differences in mean RT between pairs of levels within the factorial design. During all interference conditions, mean RT is higher at a greater task difficulty. (C) There was a significant effect of practice on response accuracy (acquisition 1<3). (D) There also was a significant effect of practice on RT (acquisition 1<3). Data displayed as mean difference ± SEM. *=p<0.05, **=p<0.01, ***=p<0.001.

### Reaction time increases as stimulus, rule and response interference increase

For mean RT data (‘Go’ trials only), the ANOVA identified a significant main effect of stimulus level (F(1,20)=222.03; p<0.001; *d*= 1.13; stimulus 1<stimulus 2) and rule level (F(2,19)=234.24; p<0.001; d= 0.40 (rule 1<rule 2); *d*= 2.65 (rule 2<rule 3); *d*= 2.70 (rule 1<rule 3)), and response level (F(1,20)=15.81; p<0.001; *d*= 0.19; response 1<response 2), as depicted in **Figure 2B**. There was a significant interaction between stimulus and rule (F(2,19)=22.67; p<0.001) and rule and response (F(2,19)=21.36; p<0.001), but not for stimulus and response (F(1,20)=0.00053; p=0.982). The threeway interaction was non-significant (F(2,19)=3.547; p=0.049).

### Practice is associated with increased accuracy and decreased reaction times

One-way ANOVA with repeated measures was performed to examine the effect of acquisition on performance accuracy and reaction time. There was a significant effect of acquisition on mean performance accuracy (F(2,40)= 12.634; p<0.001; d = 1.03; Acquisition 1 < Acquisition 3), as depicted in **Table 3 & Figure 2C**. Additionally, there was a significant effect of acquisition on mean RT (F(2,40)=11.736; p<0.001); d = 0.67; Acquisition 1 < Acquisition 3), as depicted in **Table 4 & Figure 2D**. All three acquisitions followed a trend towards faster speeds and increased performance accuracy with practice.

### Brain Activation during Task Performance

#### Rule Encoding generates activity in frontoparietal networks

A one-sample t-tests were conducted to examine the responses to rule complexity during the instruction phase (instruction level). Contrasts were weighted to display significant activation during instructions for rule level 3 relative to 1. This contrast generated one extensive continuous area of activation, which is problematic for cluster correction; consequently, the voxelwise filter was increased to p<0.0001 prior to FDR cluster correction at p<0.05. A network of brain regions was rendered at this heightened threshold, including visual association areas, precuneus and parietal cortices, putamen and anterior caudate and the superior, middle and inferior frontal gyri (**Table 5 & Figure 3Ai-iv**). In accordance with past instruction-based learning studies, activity was left-lateralised. See **Figure 3B** for visual comparison to instruction-based learning results from Hampshire et al., (2019).

**Figure 3.**
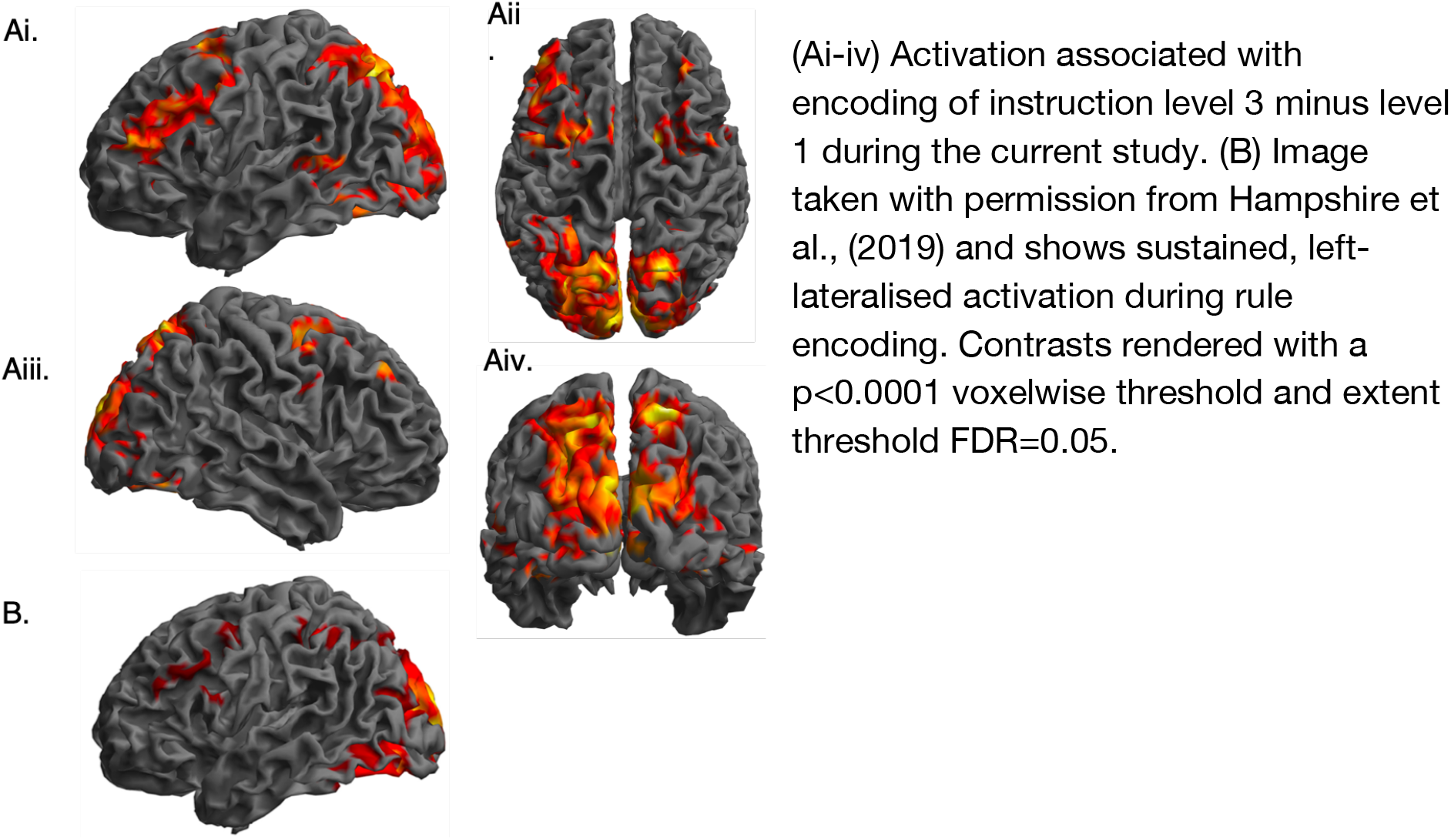
Significant activity during complex - simple rule encoding. (Ai-iv) Activation associated with encoding of instruction level 3 minus level 1 during the current study. (B) Image taken with permission from Hampshire et al., (2019) and shows sustained, left-lateralised activation during rule encoding. Contrasts rendered with a p<0.0001 voxelwise threshold and extent threshold FDR=0.05.

#### Distinct regional activation correlates of visual, rule and motor conflict

We mapped the neural correlates of increased conflict for each domain by examining the contrasts across levels separately for each domain using t-tests against 0. Contrasting high (S2) minus low (S1) visual interference trials rendered a broad pattern of occipital and parietal activity that was strongest in areas corresponding to the dorsal stream. Right-lateralised activity was also evident more focally in the frontal polar cortex and middle frontal gyrus (**Table 6 & Figure 4A**). Contrasting high (R3) minus low (R1) rule interference trials rendered widespread bilateral increases in activity within the dorsolateral prefrontal cortex (DLPFC) and frontal poles, caudate, pre-supplementary motor area (pre-SMA), anterior cingulate cortex (ACC) and superior parietal lobule (**Table 7 & Figure 4B**). Contrasting Infrequent minus frequent stop trials generated activity in areas within the mid dorsolateral prefrontal, insular, ACC and precuneus cortical areas, the anterior caudate, and proximal to left motor and somatosensory areas (**Table 8 & Figure 4C**).

**Figure 4.**
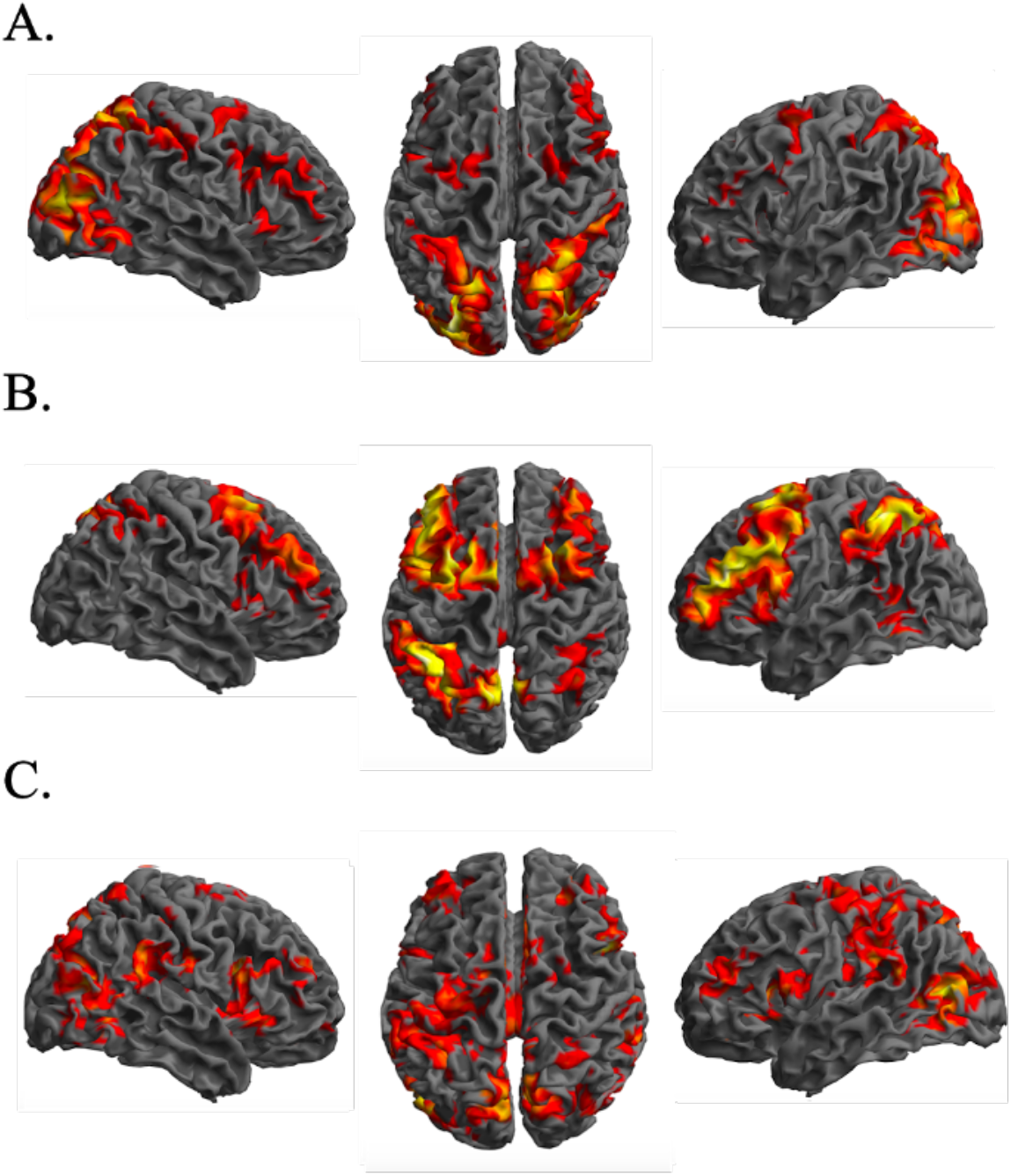
Whole-brain analysis showing regions of significant activity in response to each interference condition. (A) Visual interference. Activity spans early visual areas, spreading up through the superior parietal cortex and into frontal regions, consistent with the dorsal visual stream. (B) Rule interference. Activity spans frontal regions, including the DLPFC extending to the frontal pole and spreads through the caudate, pre-SMA, precuneus, ACC and superior parietal lobule. (C) Motor Stop Frequency. Activity spans the frontal regions as well as the insula and precuneus cortices, anterior caudate and ACC. Contrasts were rendered with a p<0.01 voxelwise threshold followed by p<0.05 FDR cluster correction.

These results appeared to indicate different patterns of brain activation when conflict was increased within the visual, rule and motor domains. However, one possibility was that these reflected the ‘imager’s fallacy’ (de Hollander et al., 2014; Henson, 2005) whereby small non-significant differences between conditions can produce visually different projections when a threshold is applied. To address this issue, we statistically analysed each contrast relative to the other two. There were areas of greater activity for visual conflict relative to rule and motor conflict (**Table 9 & Figure 5A)** spanning early visual areas along the dorsal stream to the superior parietal lobes, bilaterally. Significant activation for rule conflict relative to visual and motor frequency conflict was evident in frontal regions (**Table 10 & Figure 5B)** including left DLPFC, extending to the frontal pole, right posterior DLPFC and the parietal cortex bilaterally. Significant activation for motor frequency relative to visual and rule conflict (**Table 11 & Figure 5C)** spanned lateral occipital cortex and temporo-parietal junction (TPJ), lateral and medial temporal lobes bilaterally, anterior and posterior cingulate, left motor cortex, right anterior caudate and putamen bilaterally. Therefore, increased visual, rule and motor conflict produced significantly different patterns of network activation.

**Figure 5.**
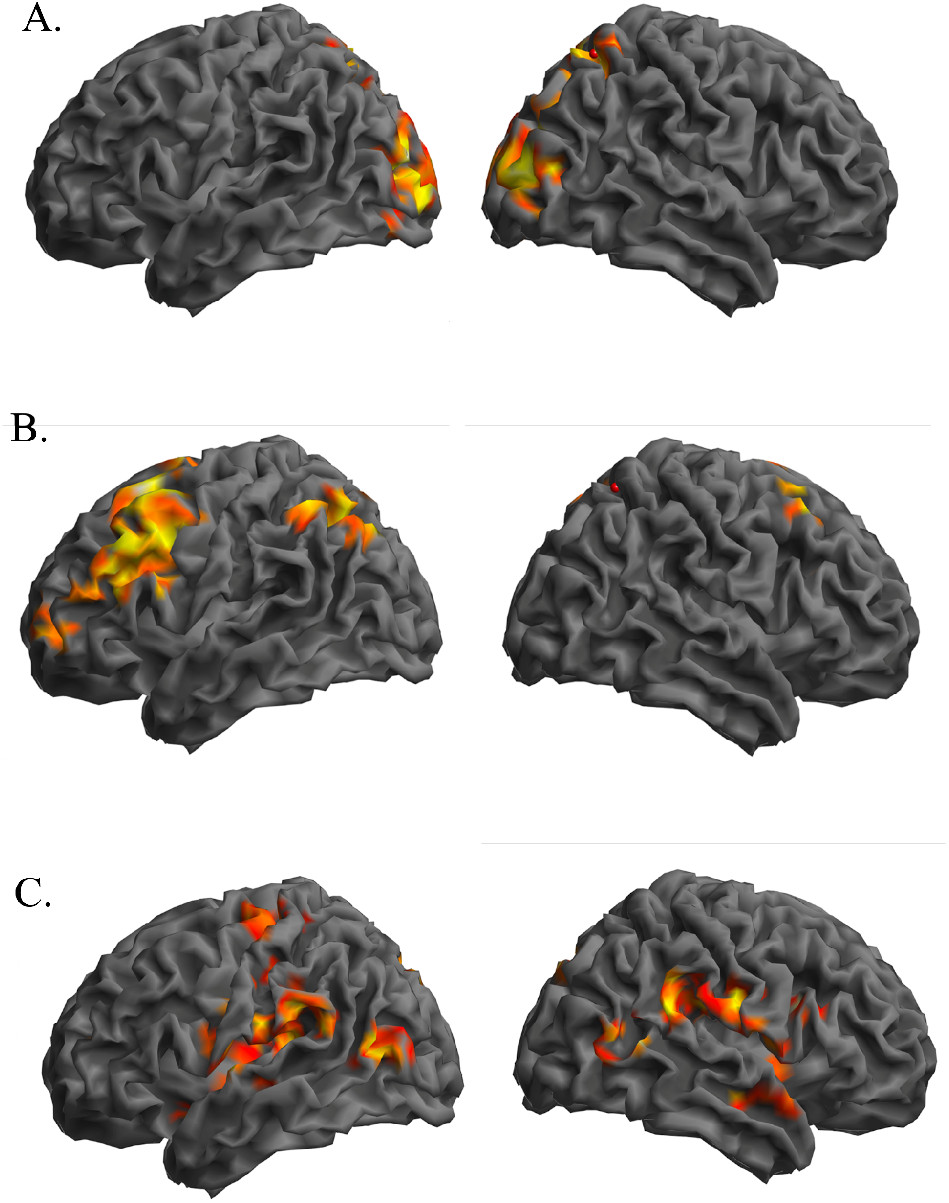
Contrasting between conflict manipulation effects. A. Visual>Rule+Motor. Activity spans early visual areas and spreads up through the superior parietal cortex. B. Rule>Visual+Motor. Activity spans frontal regions, including the left DLPFC extending to the frontal pole, right posterior DLPFC and the parietal cortex bilaterally. C. Motor Freq>Visual+Rule. Activity spans the lateral occipital cortices and temporo-parietal junction, bilaterally. Contrasts were rendered with a p<0.01 voxelwise threshold followed by p<0.05 FDR cluster correction.

### Conjunction Analysis

#### Stimulus, rule and motor conflict recruit sub-regions of Multiple Demand Cortex

Taken across all three interference conditions, the conjunction analysis rendered a network of brain regions primarily within Multiple Demand Cortex, comprising anterior insula, middle frontal gyrus (MFG) and superior frontal gyrus (SFG), bilaterally and a region within the left inferior parietal cortex (**Table 12 & Figure 6).**

**Figure 6.**
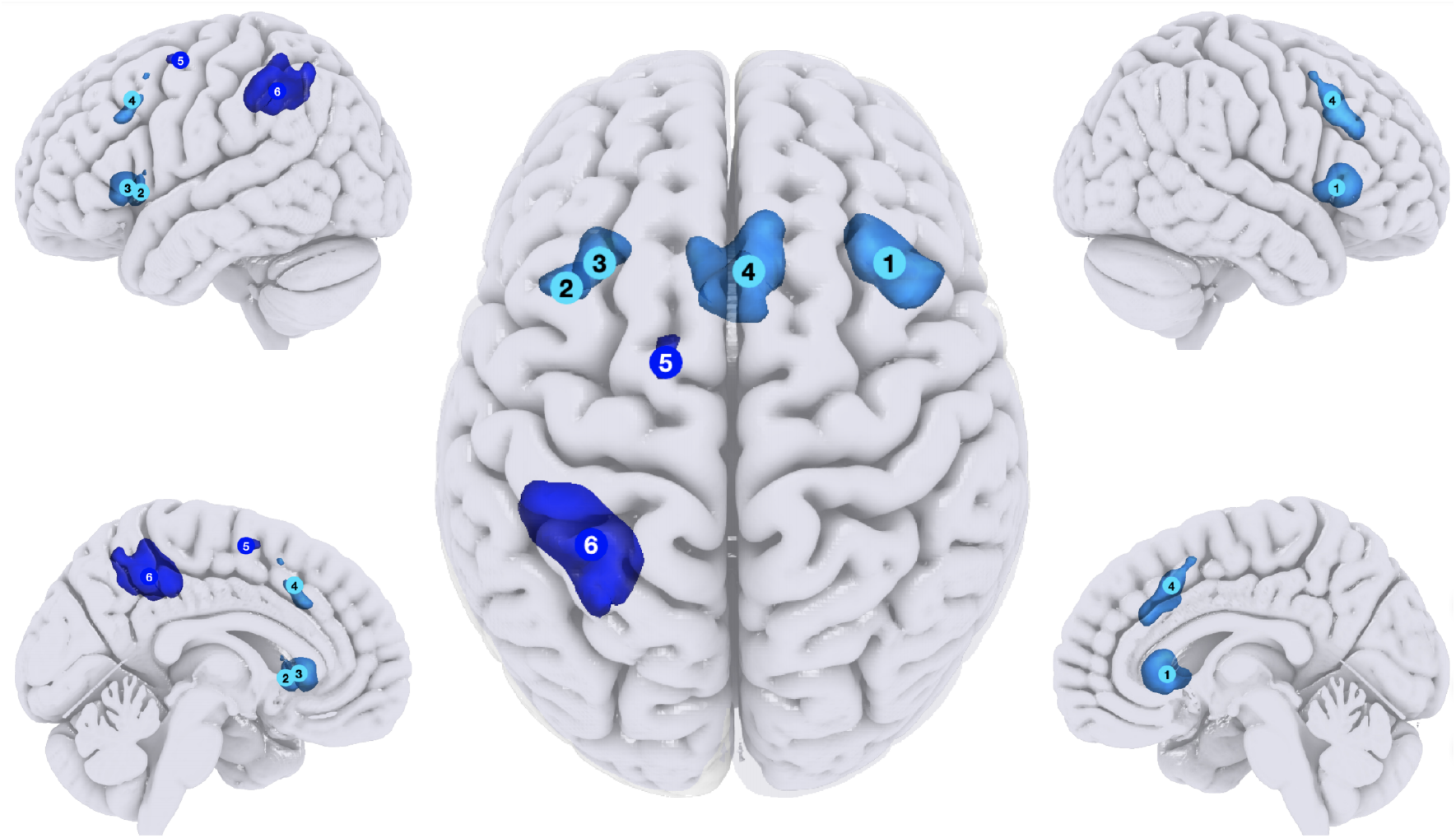
Conjunction analysis showing regions of significant activity across all three conflict conditions (Visual, Rule, Motor). Activation map depicts activity from the conjunction analysis. This primarily includes regions within MDC. Contrasts rendered at p<0.05 FDR cluster corrected.

#### Focused ROI analysis of Multiple Demand Cortex

It is hard to determine from voxelwise contrasts whether significant differences in activation between conditions are statistical or a matter of degree. Similarly, it is not possible to determine from conjunction analyses whether commonly activated areas are substantially more active for some conditions than others. Consequently, we conducted a focused analysis of interest from which MDC is comprised by parcellating an established mask (Fedorenko et al., 2013), produced by analysing activation across multiple demanding cognitive tasks, into discrete clusters using our in-house developed watershed function (**Soreq et al., 2019 & 2020**).

Repeated measures ANOVA with the factors ROI (18) and conflict domain (4) showed the expected significant ROI * condition interaction (F(34,646) =4.79; p<0.001) (**Figure 7**). There also was a significant main effect of ROI (F(17,323) =4.925; p<0.001) but not conflict domain (F(2,38) =0.637; p=0.496). Notably though, when applying liberal one tailed t-tests against 0, only one ROI showed a negative coefficient for a contrast, and only four of the eighteen ROIs were not significantly activated by all three all three types of conflict contrast, these being cerebellum area V, right lateral occipital cortex, right precentral gyrus and one of the two regions within right mid dorsolateral frontal cortex. Therefore, the dissociations between MD sub-regions primarily reflected varying degrees of sensitivity to conflict type.

**Figure 7.**
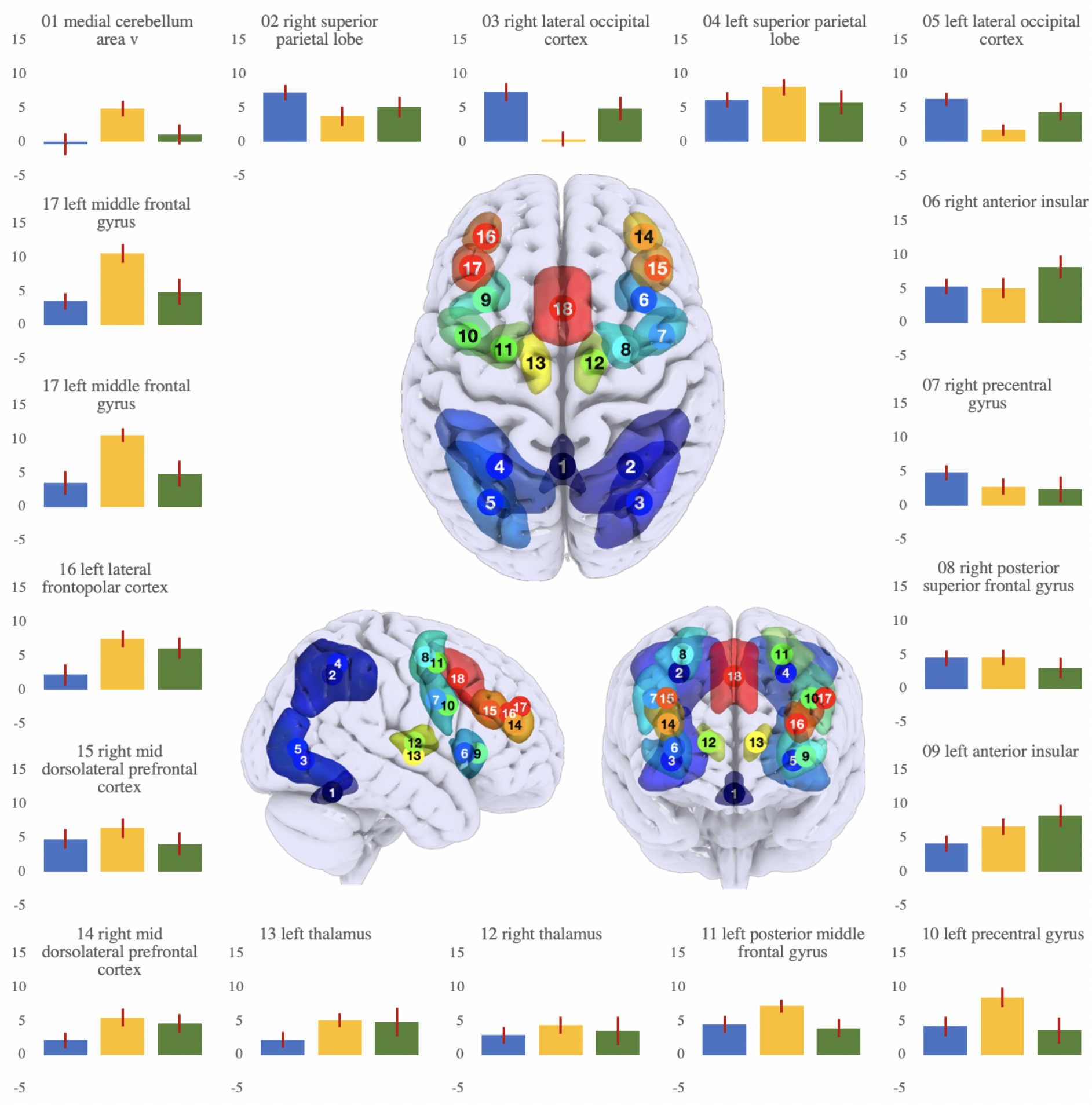
Peak beta values across 18 sub-regions of the multiple-demand cortex during rule, visual and motor frequency conflict. Parameter estimates for each MD ROI under increased visual conflict (blue), rule conflict (orange) and motor conflict (green). Y axis is in arbitrary activation units. Error bars are SEM. Note that although most ROIs were activated by all conflict manipulations, they differed substantially in their relative sensitivities.

## Discussion

Our results indicate that neither the globalist nor localist perspective on conflict resolution is entirely correct. Regarding the former, voxelwise analysis rendered significantly different patterns of activation dependent on the stage of the stimulus-response process where conflict is manipulated. These different patterns of activity were not a consequence of ‘imager’s fallacy’ (de Hollander et al., 2014; Henson, 2005), as demonstrated by direct statistical comparison between the main effect contrasts. Nor could they be explained by broader recruitment of brain regions as general difficulty increased because there was a three-way dissociation in activity, and all three manipulations (visual, rule and motor) were associated with behavioural costs during ‘Go’ trials. Furthermore, whilst these activation patterns included areas considered to have domain-specific functional roles (e.g., visual processing streams and motor cortex), they also included different sub-regions of multiple demand cortex. Considered in isolation, these observations appear to accord with the localist view that conflict of different types is processed within domain-specialised brain systems.

Counter to this interpretation, the conjunction analysis rendered areas of significant overlap between the visual, rule and motor conflict contrasts. The application of a factorial design that precisely manipulates different aspects of the same task means that it is unlikely that some other confounding process forms the basis of this observed conjunction. For example, activity related to basic visual processing and motor response, or cognitive processes such as switching and attentional orienting, are effectively subtracted out when estimating the main effects. Furthermore, significant behavioural interactions were evident within the factorial design, whereby concurrent increases in multiple domains produced non-additive costs. These latter results accord well with the notion that a common system is involved in resolving diverse conflict demands.

Notably though, it would be incorrect to conclude that the brain regions identified in the conjunction are specialised for conflict resolution. More specifically, they included anterior insula/inferior frontal operculum, anterior cingulate/ pre-supplementary motor area, inferior parietal and dorsolateral prefrontal cortex. These all are areas that fall within the classic multiple demand cortex volume (Duncan, 2001). MDC is defined by its role in diverse cognitively and attentionally demanding tasks (Duncan, 2010). Representative examples include tasks that tap processes related to conflict resolution such as response inhibition (Darda and Ramsey, 2019; Erika-Florence, 2014; Hampshire et al., 2010; Li et al., 2006), and switching (Cools et al., 2002; Daws et al., 2020; Dove et al., 2000; Hampshire and Owen, 2006), but also other processes such as target detection (Hampshire et al., 2008; Hampshire et al., 2010; Linden et al., 1999), working memory (Cole and Schneider, 2007; Nyberg et al., 2003; Soreq et al., 2019; Stiers et al., 2010), planning (Cole and Schneider, 2007) and reasoning (Crittenden et al., 2016; Hampshire et al., 2012). These latter types of task often do not have overt conflict demands.

Taken together, we believe that the above evidence accords best with a model whereby conflict resolution is an emergent property of interactions that occur across distributed networks composed of domain-specific and multiple demand brain regions, the latter of which are recruited under a broader range of contexts where additional cognitive or attentional resources are required. This simple domain specific-domain general interpretation is attractive as it can reconcile the apparent discrepancy between evidence for classic localist and globalist perspectives within a network-science framework.

Mechanistically, it seems reasonable to infer that MDC works in a top-down manner to resolve conflict between representations that occur within domains specialised systems, e.g., the visual processing streams during for visuo-perceptual conflict. Such interactions would be expected to produce increased processing load within both types of system in parallel. This view also is appealing because it accords well with models of cognitive control that are based around the principle of top-down modulation with lateral inhibition, which has been long established within the electrophysiology literature on attentional control processes (Desimone and Duncan, 1995).

However, whilst such top-down bias is likely to be part of the mechanism for conflict resolution, we argue that this view still does not fully explain the pattern of results observed here. Conflict may well occur within MDC, for example, rule conflict only activated MDC. Coding also is evident for relevant stimuli and responses within MDC as shown by human multivariate pattern analysis of imaging data (Li et al., 2007; Rao et al., 1997; Woolgar et al., 2011a; Woolgar et al., 2011b) and finer grained multi-unit analysis of the non-human primate analogues of MDC (Duncan, 2001; Freedman et al., 2001; Hoshi et al., 1998; Sigala et al., 2008).

More problematic for this view are the results of the ROI analyses, which focused on the sensitivity of MDC sub-regions to the different conflict manipulations. Each of the three interference conditions recruited almost all 18 MDC sub-regions, but to substantially varying degrees. Consequently, from a functional standpoint, it is more accurate to infer that MDC responds to conflict of different types in a coordinated but heterogeneous manner. One explanation for such multi-way dissociations could be that MDC sub-regions have more direct access to different types of information; therefore, they respond to varying levels based on the functional-anatomical locus of the conflict (Assem et al., 2020; Duncan, 2001; Duncan, 2010). Another, non-mutually exclusive, interpretation is that MDC is able to support diverse tasks in part because its sub-regions operate in different configurations.

In this respect, an interesting parallel can be drawn to our recent work investigating the functional-anatomical correlates of working memory (WM) processes (Soreq et al, 2019). Working memory studies also have previously polarised the debate between localist versus globalist perspectives. In line with the present study, we found that one-to-one mapping of specialist regions were better described as prominent features of wider, multivariate activity patterns within MDC. This led us to propose a network coding perspective whereby WM visual domains and subprocesses are classifiable by distinct functional profiles, including within MDC. Relatedly, a study by Lorenz et al, (2018) identified distinct, functional activation profiles for two networks within MDC (the dorsal frontoparietal network (FPN) and ventral FPN) within a multivariate task space.

This balance within MDC of global recruitment, and internal activation patterns that vary according to process or information content, has been observed in several other contexts, including at a fine grain for rule mapping (Woolgar et al., 2011b). Such flexibility is also evident in connectivity both with MDC, and between MDC and other brain regions (Cole et al., 2013b; Soreq et al., 2019 & 2020). As opposed to undermining the MDC hypothesis, we consider these observations to further emphasise the highly flexible nature of this global neural resource.

In conclusion, we propose that the ubiquitous process of conflict resolution is best considered as an emergent property of distributed networks, the functional-anatomical components of which sit on a continuous, not categorical, scale from domain-specialised to domain general. Brain regions that typically are labelled as MDC place towards one end of that scale; however they still have considerable functional heterogeneity, and substantial variability in their degree of domain generality, with anterior insula/inferior frontal operculum and anterior cingulate/pre-supplementary motor areas having amongst the most uniform sensitivity to different types of conflict demand. In this manner, the evidence for localist and globalist perspectives on conflict resolution, and the notion of a global resource within MDC, may be reconciled within a contemporary network science framework. Looking forward, a further challenge exists in understanding how dynamic interactions throughout these networks enable different types of conflict resolution, and whether these can be augmented in targeted ways using training or stimulation. The full factorial design reported here may also form a good basis for causal modelling and higher temporal resolution analyses of conflict resolution mechanisms.

## Materials and Methods

### Task Design

We developed a novel stimulus-response paradigm with a mixed block/event-related design that used motion-coherence, relational rule and Go/No-Go manipulations to vary conflict at different stages of the stimulus-response process (**Figure 1**). More specifically, on each trial, the participant had to discriminate the dominant direction of movement of a panel of dots, and based on a mapping rule of varying complexity, either make or withhold a motor response.

The total duration of the task was 31 minutes 12 seconds. Three independent acquisitions of the task were completed within this time. Each acquisition had a duration of 10 minutes 24 seconds. The task was run in 36 blocks, with 12 blocks completed in each of the three acquisitions, each block with a duration of 52 seconds. At the start of each block, an instruction slide with the rule and responses was shown and the participant had sixteen seconds to encode it. All instructions were presented as text. Subsequently, ten trials were presented with a fixed duration of 2.1 seconds, followed by an interstimulus-interval of 0.5 seconds. A fixation cross was displayed during the interstimulus interval. At the end of each block, there was a rest period of ten seconds before the start of the next block, enabling task-related activity to be estimated relative to a resting baseline (**Figure 1**). Participants received no feedback during the task; therefore, the mapping rules were established by a process of instruction-based learning (Cole et al., 2013a; Hampshire et al., 2016 & 2019; Ruge and Wolfenstellar, 2013).

The task parameters were manipulated in a three-dimensional full factorial design that introduced conflict for visual discrimination (2 levels), mapping rule selection (3 levels) and motor-response generation (2 levels). Each level was sampled 16 times for visual and motor factors, and 12 times for the rule factor.

Visual conflict was manipulated across two levels using dot-motion stimuli. Each stimulus consisted of 500 dots moving across a square field of view. On each trial, a larger proportion of dots moved in one direction (either up or down), with the remainder moving in the opposite direction. The proportion of dots moving in the nondominant direction was either low (0.8:0.2) or high (0.6:0.4), with the latter condition designed to produce more conflict between representations of up and down motion.

The level of conflict between alternative mapping rules was varied across three levels by manipulating the degree to which their sub-rules overlapped. In the low interference condition the direction of motion mapped uniquely to the motor response; e.g., if the dominant direction of the dots was upwards and the participant was required to make a motor response then this would be classed as a ‘Go’ trial. Alternatively, if the dominant direction was downwards and the participant was required to withhold a motor response then this would be classed as a ‘No-Go’ trial. In this condition, the dots were all coloured white (**Figure 1A**). In the medium condition, the dots were coloured either red or cyan and the mapping was dependent on the conjunction of direction and colour (**Table 1; Figure 1B**). This makes it harder to resolve between the alternative mapping rules given the stimulus features considered individually map to both response conditions. The dot conditions (the dominant direction of motion and the direction-colour combinations) were pseudorandomised to ensure a given stimulus condition or required response occurred no more than three times in succession. The high rule-conflict condition was the same as the medium condition but with the addition of a temporal component such that the conjunction of stimulus features also mapped to both response conditions (**Table 2; Figure 1B**). Specifically, as opposed to the colour-motion combinations mapping to a specific response, participants were required to either repeat the response they gave during the previous trial (t-1) or select the alternate response (i.e. Go if the previous trial was No-Go). At this harder level, for the first trial of each block, participants were required to respond as per the medium rule level (**Table 1**).

To examine the level of motor conflict, the frequency of Go vs No-Go responses was varied from either 3:7 or 7:3. This was implemented based on the hypothesis that omission of a response in the No-Go condition should be more difficult if the motor responses are frequent and therefore prepotent (Nieuwenhuis et al., 2003, Hampshire and Sharp, 2015).

Prior to data collection, participants read a written protocol and were given the opportunity to ask questions about the task, to ensure they fully understood the requirements. Following this, participants completed three shortened practice acquisitions outside of the scanner, each containing two or three blocks. Each shortened practice acquisition gave an example of one task dimension changing over all levels of interference, whilst the levels of the other two dimensions remained constant, and at the simplest level, throughout. Following this, participants completed two full practice acquisitions of the task before entering the scanner. The total time for the practice phase was 26 minutes 52 seconds.

### Behavioural analysis

The task was programmed using Psychtoolbox for MATLAB (Brainard, 1997). Three variants of the task paradigm were tested prior to arriving at the design reported here. This was to (1) identify the optimal amount of practice to produce consistent performance across runs, (2) ensure that the task produced the desired behavioural interference effects, whilst (3) confirming that participants could perform the more difficult task conditions above chance level. Mean reaction time (RT) (calculated as the time between trial onset and button response during ‘Go’ trials) and response accuracy (RA) (calculated as the number of correct responses within 10 trials) were computed using MATLAB (MathWorks). A factorial analysis of variance (ANOVA) was performed, using RT and RA, to test for a main effect of stimulus level, rule level or response level and for interaction effects between factors on both measures. A oneway ANOVA with repeated measures and Tukey HSD post-hoc test were performed to examine if there was an effect of run on mean RT and/or RA and therefore to determine the optimum length of the practice phase. Effect size for the pairwise comparisons was calculated using η2 (η2 = between groups sum of squares / total sum of squares) and compared to Cohen’s *d.*

### fMRI acquisition and preprocessing

21 individuals (14=female, mean age=23.10 yrs, SD=0.77, range=21-25 yrs) performed the interference task for 31 minutes and 12 seconds. Scanning was undertaken on a 3T Siemens Verio (Siemens, Erlangen, Germany), using a 32-channel head coil. Functional images were collected using the following parameters: T2*-weighted gradient-echo, echoplanar imaging (EPI) sequence, TR=2s, TE=30ms, voxel size=3mm^3^, FA=80°, FOV=192×192×105 mm, 35 slices, GRAPPA=2. A T1-weighted image was also acquired using an MPRAGE sequence, TR=2.3s, TE=2.98ms, TI=900ms, voxel size=1mm^3^, FA=9°, FOV=256×256 mm, 256×256 matrix, 160 slices, GRAPPA=2. FMRI data were preprocessed using FSL (Jenkinson et al., 2012) and SPM12 (Ashburner et al., 2014) tools. In brief, images were motion corrected using MCFLIRT (Jenkinson et al., 2002) and coregistered to the BET (Smith, 2002) T1-image using boundary-based registration (BBR) (Greve and Fischl, 2009). We then used DARTEL to generate a study template and to calculate the structural data warps to MNI space and then applied these to normalise the fMRI images. Normalised images were smoothed with a 5mm full-width at half maximum Gaussian kernel.

### Image Analysis

Single subject task-evoked brain activation patterns were estimated by general linear modelling (GLM) in SPM12. 36 predictor variables were generated from the relevant task events via the convolution of task onset timings and durations with the haemodynamic response function (HRF), providing estimates of event related brain activation per unit time. These included instructions for blocks 1-12, ‘Go’ trials for blocks 1-12 and ‘No-Go’ trials for blocks 1-12. Rotations and translations in the X, Y and Z planes, calculated during motion correction, were included as six nuisance variables.

Contrast images were generated for group level analysis. These were: instruction difficulty (all hard minus easy instructions), rule complexity (all complex minus simple mapping trials (R3-R1)), visual discriminability (all hard minus easy stimulus level trials (S2-S1)) and motor response frequency (all infrequent minus all frequent ‘No-Go’ trials).

Contrast images from the individual subject models were examined at the group level using one sample t-tests, one-way ANOVAs and factorial designs in SPM12. Group-level analyses were FDR cluster correction for the whole-brain volume at p<0.05 after applying a voxelwise filter of p<0.01. Activation labels were determined from MNI coordinates using the Harvard-Oxford structural atlas in FSLeyes (Jenkinson et al., 2012). Common regions of activation in response to all three interference conditions were determined using conjunction analysis with whole brain cluster correction at FDR p<0.05. Statistical maps were thresholded voxelwise at p<0.01 uncorrected prior to conjunction.

For the Region-of-interest (ROI) analysis, the statistical Multiple Demand Cortex volume published by Fedorenko et al. (2013) was used to define both MDC (i.e. task active) and DMN (i.e. task negative) ROI sets. Using our in-house fusion-watershed toolbox (Soreq et al., 2019 & 2020 Nat Comms) both functional activation maps were segmented into discrete clusters. A 150 voxels threshold was applied with a 6mm neighbourhood radius around the local peak maxima. At the end of the clustering stage small ROI’s (<50 voxels) were merged with bigger neighbours. Mean contrast values for all voxels within each ROI were averaged and extracted for each subject. These ROI activation estimates were analysed using repeated measures ANOVA in SPSS.

## Supplementary Tables

**Table 3.** Paired t-tests to determine the effect of run on response accuracy

|  |  |  |  | 95% Confidence Interval of the Difference |  |  |  |  |
| --- | --- | --- | --- | --- | --- | --- | --- | --- |
| Run | Mean | Std. Deviation | Std. Error Mean | Lower | Upper | t | df | Sig. (2-tailed) |
| 1 | -.24206 | .46837 | .10221 | -.45526 | -.02886 | -2.368 | 20 | .028 |
| 2 | -.56745 | .58844 | .12841 | -.83530 | -.29960 | -4.419 | 20 | .000 |
| 3 | -.32539 | .49293 | .10757 | -.54977 | -.10101 | -3.025 | 20 | .007 |

**Table 4.** Paired t-tests to determine the effect of run on response time

|  |  |  |  | 95% Confidence Interval of the Difference |  |  |  |  |
| --- | --- | --- | --- | --- | --- | --- | --- | --- |
| Run | Mean | Std. Deviation | Std. Error Mean | Lower | Upper | t | df | Sig. (2-tailed) |
| 1 | .01694 | .07716 | .01684 | -.01818 | .05206 | 1.006 | 20 | .326 |
| 2 | .07218 | .07004 | .01528 | .04030 | .10406 | 4.722 | 20 | .000 |
| 3 | .05524 | .06662 | .01454 | .02492 | .08557 | 3.800 | 20 | .001 |

**Table 5.**
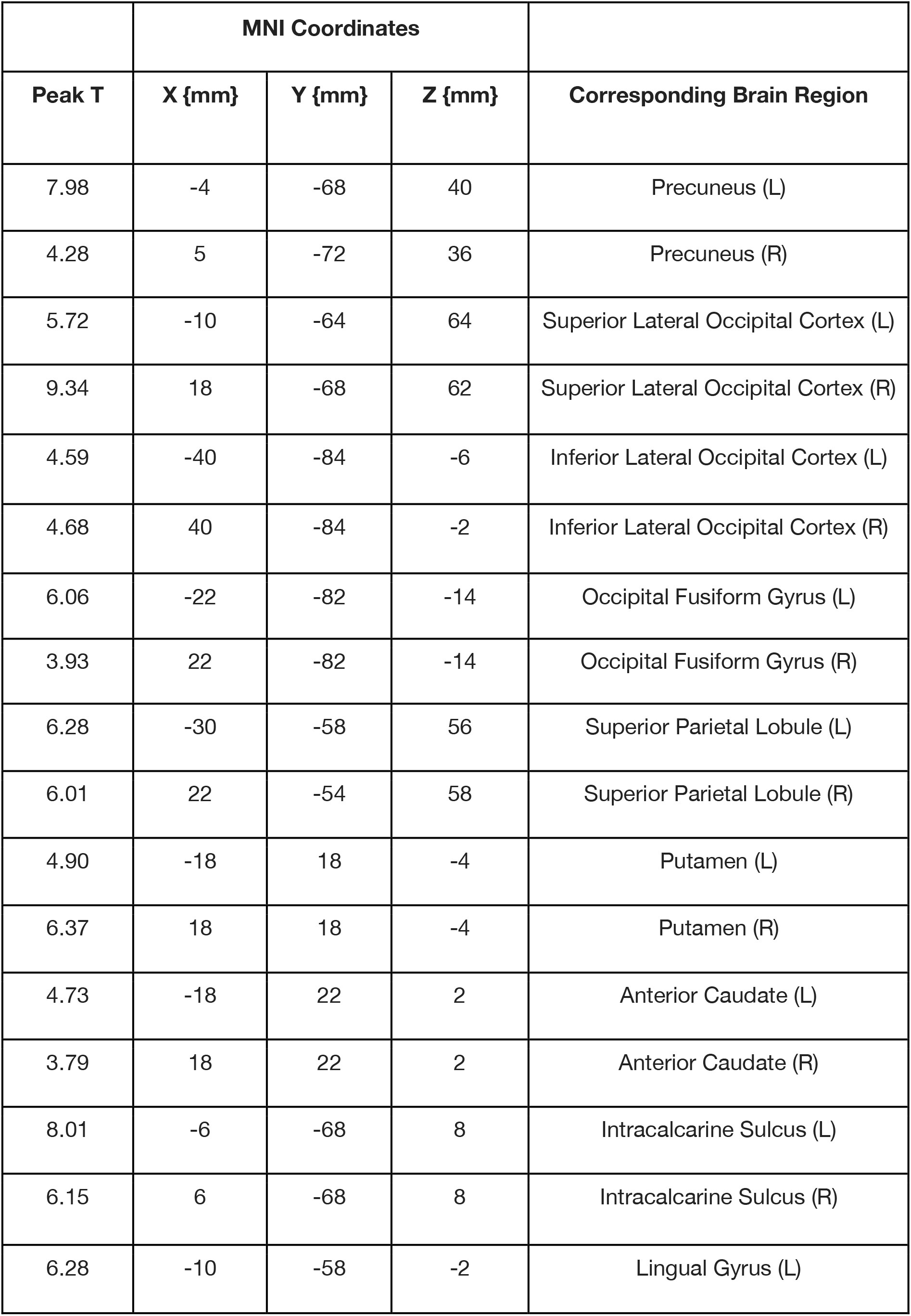
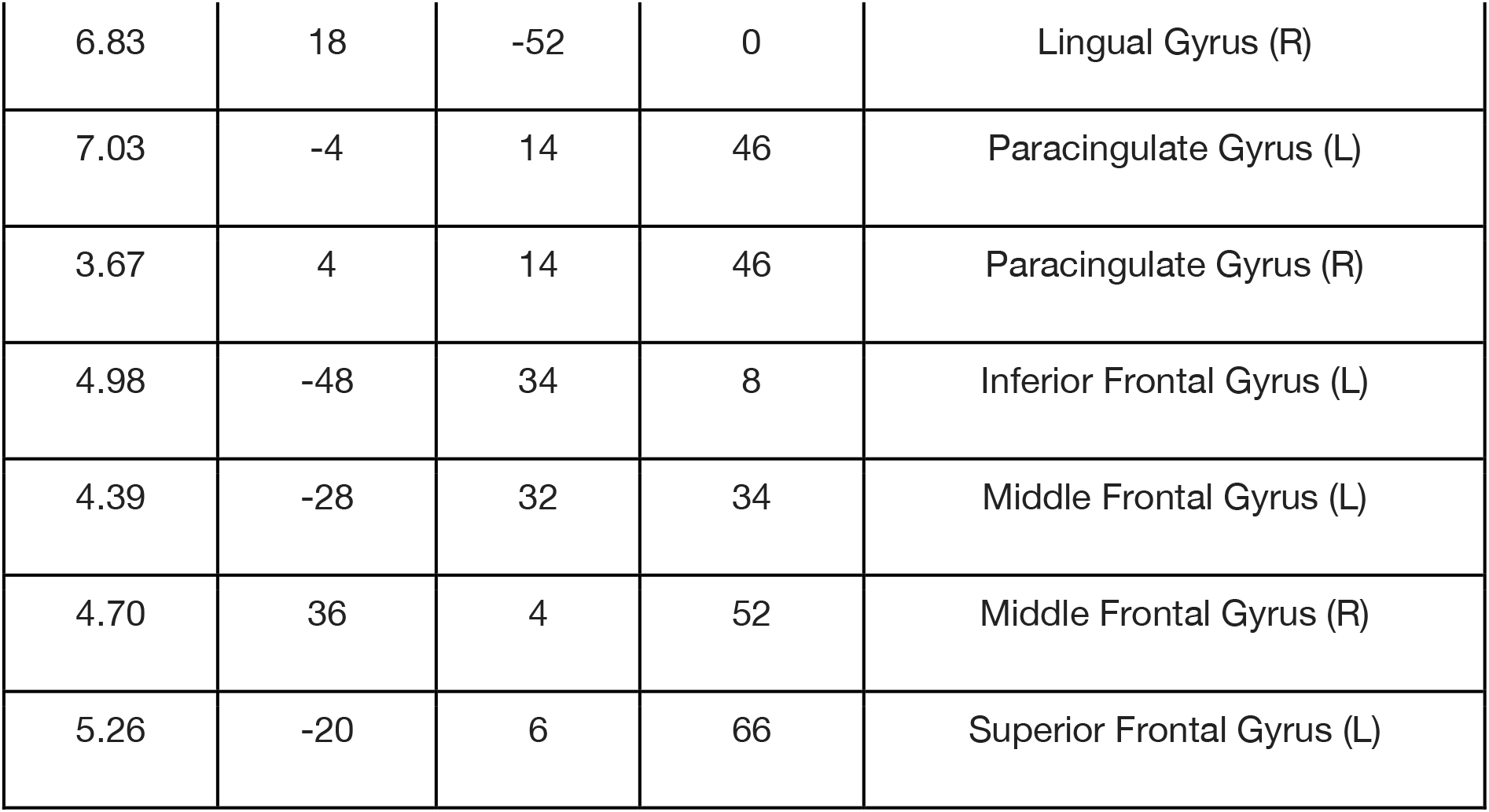
Peak values, MNI coordinates and corresponding brain regions for rule encoding.

|  | **MNI Coordinates** |  |  |  |
| --- | --- | --- | --- | --- |
| **Peak T** | **X {mm}** | **Y {mm}** | **Z {mm}** | **Corresponding Brain Region** |
| 7.98 | -4 | -68 | 40 | Precuneus (L) |
| 4.28 | 5 | -72 | 36 | Precuneus (R) |
| 5.72 | -10 | -64 | 64 | Superior Lateral Occipital Cortex (L) |
| 9.34 | 18 | -68 | 62 | Superior Lateral Occipital Cortex (R) |
| 4.59 | -40 | -84 | -6 | Inferior Lateral Occipital Cortex (L) |
| 4.68 | 40 | -84 | -2 | Inferior Lateral Occipital Cortex (R) |
| 6.06 | -22 | -82 | -14 | Occipital Fusiform Gyrus (L) |
| 3.93 | 22 | -82 | -14 | Occipital Fusiform Gyrus (R) |
| 6.28 | -30 | -58 | 56 | Superior Parietal Lobule (L) |
| 6.01 | 22 | -54 | 58 | Superior Parietal Lobule (R) |
| 4.90 | -18 | 18 | -4 | Putamen (L) |
| 6.37 | 18 | 18 | -4 | Putamen (R) |
| 4.73 | -18 | 22 | 2 | Anterior Caudate (L) |
| 3.79 | 18 | 22 | 2 | Anterior Caudate (R) |
| 8.01 | -6 | -68 | 8 | Intracalcarine Sulcus (L) |
| 6.15 | 6 | -68 | 8 | Intracalcarine Sulcus (R) |
| 6.28 | -10 | -58 | -2 | Lingual Gyrus (L) |
| 6.83 | 18 | -52 | 0 | Lingual Gyrus (R) |
| 7.03 | -4 | 14 | 46 | Paracingulate Gyrus (L) |
| 3.67 | 4 | 14 | 46 | Paracingulate Gyrus (R) |
| 4.98 | -48 | 34 | 8 | Inferior Frontal Gyrus (L) |
| 4.39 | -28 | 32 | 34 | Middle Frontal Gyrus (L) |
| 4.70 | 36 | 4 | 52 | Middle Frontal Gyrus (R) |
| 5.26 | -20 | 6 | 66 | Superior Frontal Gyrus (L) |

**Table 6.**
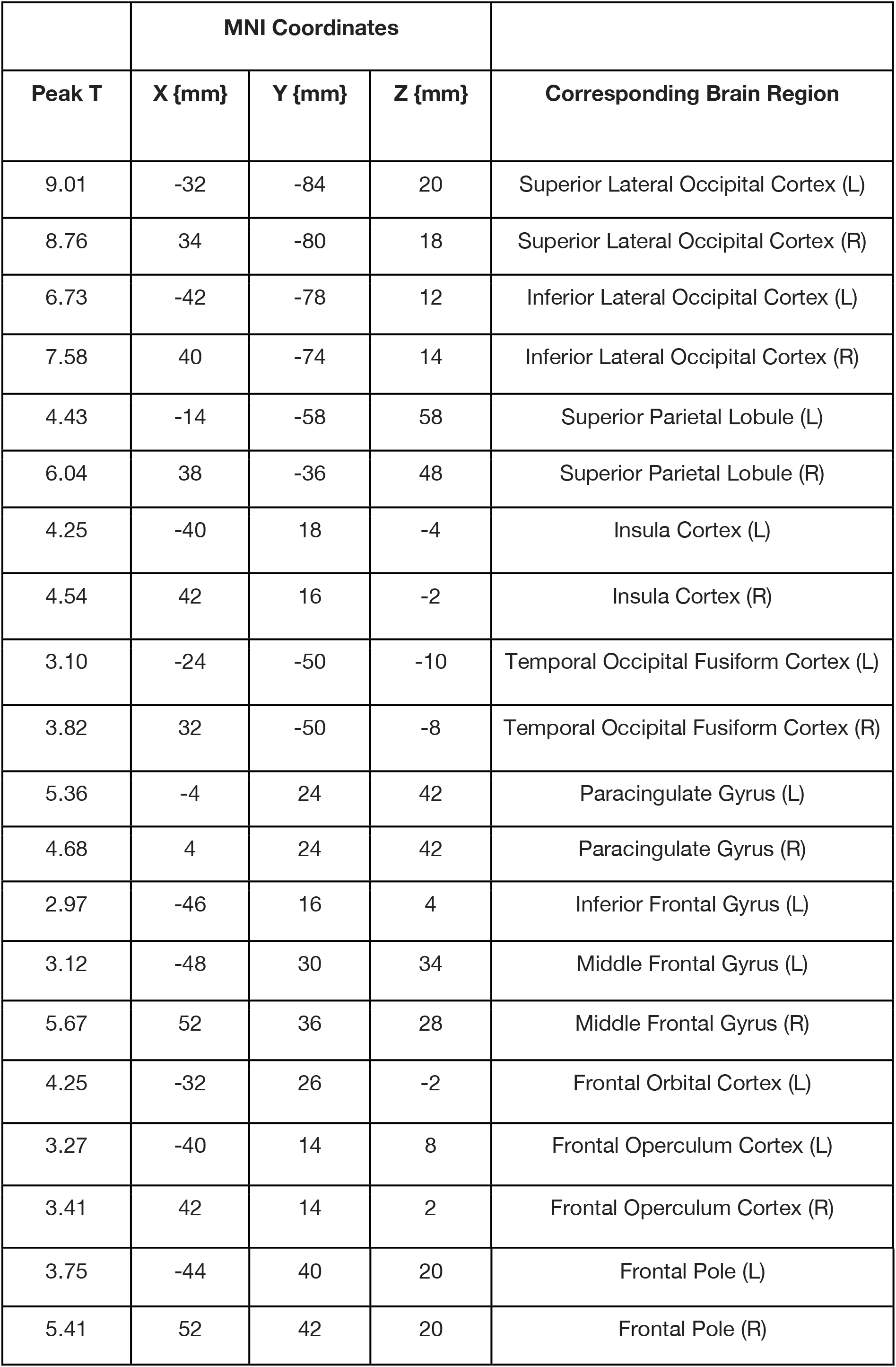
Peak values, MNI coordinates and corresponding brain regions for visual difficulty.

|  | **MNI Coordinates** |  |  |  |
| --- | --- | --- | --- | --- |
| **Peak T** | **X {mm}** | **Y {mm}** | **Z {mm}** | **Corresponding Brain Region** |
| 9.01 | -32 | -84 | 20 | Superior Lateral Occipital Cortex (L) |
| 8.76 | 34 | -80 | 18 | Superior Lateral Occipital Cortex (R) |
| 6.73 | -42 | -78 | 12 | Inferior Lateral Occipital Cortex (L) |
| 7.58 | 40 | -74 | 14 | Inferior Lateral Occipital Cortex (R) |
| 4.43 | -14 | -58 | 58 | Superior Parietal Lobule (L) |
| 6.04 | 38 | -36 | 48 | Superior Parietal Lobule (R) |
| 4.25 | -40 | 18 | -4 | Insula Cortex (L) |
| 4.54 | 42 | 16 | -2 | Insula Cortex (R) |
| 3.10 | -24 | -50 | -10 | Temporal Occipital Fusiform Cortex (L) |
| 3.82 | 32 | -50 | -8 | Temporal Occipital Fusiform Cortex (R) |
| 5.36 | -4 | 24 | 42 | Paracingulate Gyrus (L) |
| 4.68 | 4 | 24 | 42 | Paracingulate Gyrus (R) |
| 2.97 | -46 | 16 | 4 | Inferior Frontal Gyrus (L) |
| 3.12 | -48 | 30 | 34 | Middle Frontal Gyrus (L) |
| 5.67 | 52 | 36 | 28 | Middle Frontal Gyrus (R) |
| 4.25 | -32 | 26 | -2 | Frontal Orbital Cortex (L) |
| 3.27 | -40 | 14 | 8 | Frontal Operculum Cortex (L) |
| 3.41 | 42 | 14 | 2 | Frontal Operculum Cortex (R) |
| 3.75 | -44 | 40 | 20 | Frontal Pole (L) |
| 5.41 | 52 | 42 | 20 | Frontal Pole (R) |

**Table 7.** Peak values, MNI coordinates and corresponding brain regions for rule difficulty.

|  | **MNI Coordinates** |  |  |  |
| --- | --- | --- | --- | --- |
| **Peak T** | **X {mm}** | **Y {mm}** | **Z {mm}** | **Corresponding Brain Region** |
| 6.58 | -38 | -60 | 50 | Superior Lateral Occipital Cortex (L) |
| 4.68 | 38 | -60 | 50 | Superior Lateral Occipital Cortex (R) |
| 8.33 | -28 | -54 | 46 | Superior Parietal Lobule (L) |
| 4.10 | 38 | -48 | 54 | Superior Parietal Lobule (R) |
| 8.41 | -44 | -46 | 56 | Posterior Supramarginal Gyrus (L) |
| 3.47 | 48 | -44 | 58 | Posterior Supramarginal Gyrus (R) |
| 7.13 | -4 | 4 | 66 | Supplementary Motor Cortex (L) |
| 8.44 | -4 | -66 | 56 | Precuneus Cortex (L) |
| 7.38 | 1 | -64 | 46 | Precuneus Cortex (R) |
| 9.28 | -2 | 16 | 48 | Paracingulate Gyrus (L) |
| 5.15 | 10 | 14 | 46 | Paracingulate Gyrus (R) |
| 5.68 | -30 | 24 | 2 | Insula Cortex (L) |
| 4.68 | -18 | 14 | 10 | Caudate (L) |
| 4.50 | 18 | 22 | 4 | Caudate (R) |
| 3.35 | -10 | 22 | 28 | Anterior Cingulate Cortex (L) |
| 3.84 | 6 | 28 | 28 | Anterior Cingulate Cortex (R) |
| 6.09 | -44 | 6 | 22 | Central Opercular Cortex (L) |
| 4.11 | -54 | -58 | -10 | Inferior Temporal Gyrus (L) |
| 4.12 | -58 | -56 | 4 | Middle Temporal Gyrus (L) |
| 5.06 | 36 | 26 | 0 | Frontal Orbital Cortex (R) |
| 4.90 | 46 | 18 | 0 | Frontal Operculum Cortex (R) |
| 4.78 | -52 | 18 | 0 | Inferior Frontal Gyrus (L) |
| 4.78 | -38 | 28 | 46 | Middle Frontal Gyrus (L) |
| 4.73 | 44 | 28 | 32 | Middle Frontal Gyrus (R) |
| 9.10 | -26 | 14 | 60 | Superior Frontal Gyrus (L) |
| 6.16 | 24 | 10 | 66 | Superior Frontal Gyrus (R) |
| 8.81 | -44 | 40 | 18 | Frontal Pole (L) |
| 3.37 | 36 | 42 | 8 | Frontal Pole (R) |

**Table 8.** Peak values, MNI coordinates and corresponding brain regions for motor response frequency.

| Peak T | MNI Coordinates |  |  | Corresponding Brain Region |
| --- | --- | --- | --- | --- |
|  | X {mm} | Y {mm} | Z {mm} |  |
| 4.26 | -48 | -76 | -2 | Inferior Lateral Occipital Cortex (L) |
| 4.06 | 52 | -68 | 2 | Inferior Lateral Occipital Cortex (R) |
| 3.79 | -26 | -84 | 32 | Superior Lateral Occipital Cortex (L) |
| 3.81 | 34 | -80 | 38 | Superior Lateral Occipital Cortex (R) |
| 4.22 | -34 | -48 | 50 | Superior Parietal Lobule (L) |
| 3.76 | 26 | -54 | 64 | Superior Parietal Lobule (R) |
| 5.20 | -20 | -58 | 6 | Precuneus Cortex (L) |
| 5.19 | 18 | -58 | 8 | Precuneus Cortex (R) |
| 5.38 | -14 | 20 | 0 | Caudate (L) |
| 7.54 | 8 | 18 | 2 | Caudate (R) |
| 7.05 | 30 | -48 | -8 | Temporal Occipital Fusiform Cortex (R) |
| 5.89 | -26 | -54 | 6 | Lingual Gyrus (L) |
| 4.09 | 22 | -52 | -2 | Lingual Gyrus (R) |
| 4.28 | -2 | -34 | 26 | Posterior Cingulate Gyrus (L) |
| 4.28 | 2 | -32 | 28 | Posterior Cingulate Gyrus (R) |
| 4.08 | -16 | -6 | 62 | Superior Frontal Gyrus (L) |
| 3.32 | 24 | -2 | 66 | Superior Frontal Gyrus (R) |
| 3.83 | -28 | 46 | 12 | Frontal Pole (L) |
| 2.96 | 22 | 60 | -6 | Frontal Pole (R) |

**Table 9.** Peak values, MNI coordinates and corresponding brain regions for visual discriminability relative to the rule and motor dimensions.

|  | MNI Coordinates |  |  |  |
| --- | --- | --- | --- | --- |
| Peak T | X {mm} | Y {mm} | Z {mm} | Corresponding Brain Region |
| 4.24 | -18 | -92 | -2 | Occipital Pole (L) |
| 4.72 | 28 | -94 | 14 | Occipital Pole (R) |
| 5.48 | -32 | -90 | 18 | Superior Lateral Occipital Cortex (L) |
| 4.81 | 24 | -64 | 38 | Superior Lateral Occipital Cortex (R) |
| 4.64 | -20 | -60 | 58 | Superior Parietal (L) |
| 4.24 | 22 | -58 | 58 | Superior Parietal (R) |

**Table 10.** Peak values, MNI coordinates and corresponding brain regions for rule difficulty relative to the visual and motor dimensions.

|  | **MNI Coordinates** |  |  |  |
| --- | --- | --- | --- | --- |
| **Peak T** | **X {mm}** | **Y {mm}** | **Z {mm}** | **Corresponding Brain Region** |
| 3.47 | 0 | -60 | -4 | Cerebellum (R) |
| 4.44 | -48 | -50 | 54 | Parietal Cortex (L) |
| 4.3 | -2 | 4 | 60 | SMA (L) |
| 5.09 | -4 | -64 | 52 | Precuneus (L) |
| 3.72 | -18 | 58 | 8 | Lateral Orbitofrontal Cortex (L) |
| 3.61 | -46 | 36 | 24 | Mid dorsolateral PFC (L) |
| 4.89 | -42 | 26 | 38 | Posterior Middle Frontal Gyrus (L) |
| 6.16 | -26 | 14 | 60 | Posterior Superior Frontal Cortex (L) |
| 4.23 | 28 | 16 | 46 | Posterior Superior Frontal Cortex (R) |

**Table 11.**
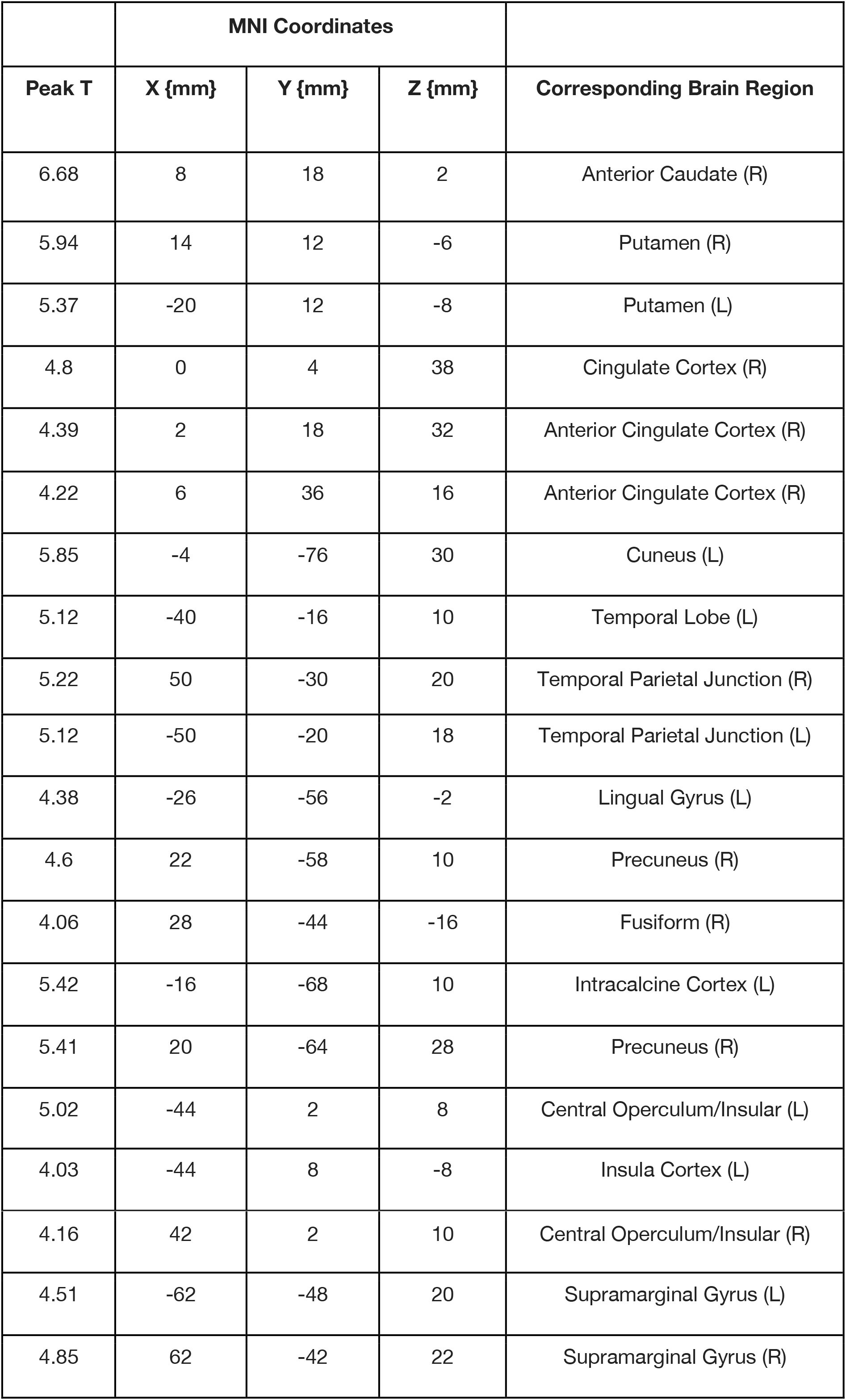
Peak values, MNI coordinates and corresponding brain regions for motor response frequency relative to the visual and rule dimensions.

**Table 12.** Peak values, MNI coordinates and corresponding brain regions for the conjunction of average activity across all interference conditions.

|  | MNI Coordinates |  |  |  |
| --- | --- | --- | --- | --- |
| Peak T | X {mm} | Y {mm} | Z {mm} | Corresponding Brain Region |
| 3.99 | -32 | -59 | 60 | Superior Occipital Cortex (L) |
| 6.86 | -25 | -54 | 46 | Superior Parietal Lobule (L) |
| 5.74 | -45 | -43 | 46 | Supramarginal Gyrus (L) |
| 4.64 | 8 | 34 | 30 | Paracingulate Gyrus (R) |
| 6.31 | -6 | 19 | 46 | Paracingulate Gyrus (L) |
| 4.95 | 30 | 20 | 10 | Insula Cortex (R) |
| 5.13 | -38 | 16 | -1 | Insula Cortex (L) |
| 3.53 | -48 | -30 | 47 | PostCentral Gyrus (L) |
| 4.94 | 8 | 31 | 23 | Anterior Cingulate Gyrus (R) |
| 5.85 | 45 | 20 | 2 | Frontal Operculum (L) |
| 5.28 | -36 | 18 | 6 | Frontal Operculum (R) |
| 4.69 | -46 | 18 | 6 | Inferior Frontal Gyrus (L) |
| 4.05 | 52 | 18 | 28 | Inferior Frontal Gyrus (R) |
| 3.79 | 49 | 20 | 44 | Middle Frontal Gyrus (R) |
| 6.14 | 2 | 16 | 59 | Superior Frontal Gyrus (R) |
| 4.25 | 40 | 36 | 22 | Frontal Pole (R) |

## References

Akçay, Ç. and Hazeltine, E., (2011). Domain-specific conflict adaptation without feature repetitions. Psychonomic Bulletin& Review, 18(3), pp.505–511.

Ambrosini, E. and Vallesi, A., (2017). Domain-general Stroop performance and hemispheric asymmetries: a resting-state EEG study. Journal of cognitive neuroscience, 29(5), pp.769779.

Ashburner, J., Barnes, G., Chen, C., Daunizeau, J., Flandin, G., Friston, K., Kiebel, S., Kilner, J., Litvak, V., Moran, R. and Penny, W., (2014). SPM12 manual. Wellcome Trust Centre for Neuroimaging, London, UK, p.2464.

Assem, M., Blank, I.A., Mineroff, Z., Ademoglu, A. and Fedorenko, E., (2020). Activity in the fronto-parietal multiple-demand network is robustly associated with individual differences in working memory and fluid intelligence. bioRxiv, p.110270.

Botvinick, M.M., Braver, T.S., Barch, D.M., Carter, C.S. and Cohen, J.D., (2001). Conflict monitoring and cognitive control. Psychological review, 108(3), p.624.

Brainard, D.H., (1997). The psychophysics toolbox. Spatial vision, 10(4), pp.433–436.

Braver, T.S., Paxton, J.L., Locke, H.S. and Barch, D.M., (2009). Flexible neural mechanisms of cognitive control within human prefrontal cortex. Proceedings of the National Academy of Sciences, 106(18), pp.7351–7356.

Cole, M.W., Laurent, P. and Stocco, A., (2013a). Rapid instructed task learning: A new window into the human brain’s unique capacity for flexible cognitive control. Cognitive, Affective,& Behavioral Neuroscience, 13(1), pp.1–22.

Cole, M.W., Reynolds, J.R., Power, J.D., Repovs, G., Anticevic, A. and Braver, T.S., (2013b). Multi-task connectivity reveals flexible hubs for adaptive task control. Nature neuroscience, 16(9), pp.1348–1355.

Cole, M.W. and Schneider, W., (2007). The cognitive control network: integrated cortical regions with dissociable functions. Neuroimage, 37(1), pp.343–360.

Cools, R., Clark, L., Owen, A.M. and Robbins, T.W., (2002). Defining the neural mechanisms of probabilistic reversal learning using event-related functional magnetic resonance imaging. Journal of Neuroscience, 22(11), pp.4563–4567.

Crittenden, B.M., Mitchell, D.J. and Duncan, J., (2016). Task encoding across the multiple demand cortex is consistent with a frontoparietal and cingulo-opercular dual networks distinction. Journal of Neuroscience, 36(23), pp.6147–6155.

Darda, K.M. and Ramsey, R., (2019). The inhibition of automatic imitation: a meta-analysis and synthesis of fMRI studies. Neuroimage, 197, pp.320–329.

Daws, R.E., Soreq, E., Li, Y., Sandrone, S. and Hampshire, A., (2020). Contrasting hierarchical and multiple-demand accounts of frontal lobe functional organisation during task-switching. bioRxiv.

de Hollander, G., Wagenmakers, E.J., Waldorp, L. and Forstmann, B., (2014). An antidote to the imager’s fallacy, or how to identify brain areas that are in limbo. PLoS One, 9(12), p.e115700.

Desimone, R. and Duncan, J., (1995). Neural mechanisms of selective visual attention. Annual review of neuroscience, 18(1), pp.193–222.

Dosenbach, N.U., Fair, D.A., Cohen, A.L., Schlaggar, B.L. and Petersen, S.E., (2008). A dual-networks architecture of top-down control. Trends in cognitive sciences, 12(3), pp.99–105.

Dove, A., Pollmann, S., Schubert, T., Wiggins, C.J. and Von Cramon, D.Y., (2000). Prefrontal cortex activation in task switching: an event-related fMRI study. Cognitive brain research, 9(1), pp.103–109.

Duncan, J., (2001). An adaptive coding model of neural function in prefrontal cortex. Nature reviews neuroscience, 2(11), pp.820–829.

Duncan, J., (2010). The multiple-demand (MD) system of the primate brain: mental programs for intelligent behaviour. Trends in cognitive sciences, 14(4), pp.172–179.

Egner, T., (2008). Multiple conflict-driven control mechanisms in the human brain. Trends in cognitive sciences, 12(10), pp.374–380.

Egner, T., Delano, M. and Hirsch, J., (2007). Separate conflict-specific cognitive control mechanisms in the human brain. neuroimage, 35(2), pp.940–948.

Erika-Florence, M., Leech, R. and Hampshire, A., (2014). A functional network perspective on response inhibition and attentional control. Nature communications, 5(1), pp.1–12.

Fan, J., Flombaum, J.I., McCandliss, B.D., Thomas, K.M. and Posner, M.I., (2003). Cognitive and brain consequences of conflict. Neuroimage, 18(1), pp.42–57.

Fedorenko, E., Duncan, J. and Kanwisher, N., (2013). Broad domain generality in focal regions of frontal and parietal cortex. Proceedings of the National Academy of Sciences, 110(41), pp.16616–16621.

Freedman, D.J., Riesenhuber, M., Poggio, T. and Miller, E.K., (2001). Categorical representation of visual stimuli in the primate prefrontal cortex. Science, 291(5502), pp.312–316.

Freitas, A.L., Bahar, M., Yang, S. and Banai, R., (2007). Contextual adjustments in cognitive control across tasks. Psychological Science, 18(12), pp.1040–1043.

Greve, D.N. and Fischl, B., (2009). Accurate and robust brain image alignment using boundary-based registration. Neuroimage, 48(1), pp.63–72.

Hampshire, A., Chamberlain, S.R., Monti, M.M., Duncan, J. and Owen, A.M., (2010). The role of the right inferior frontal gyrus: inhibition and attentional control. Neuroimage, 50(3), pp.1313–1319.

Hampshire, A., Daws, R.E., Neves, I.D., Soreq, E., Sandrone, S. and Violante, I.R., (2019). Probing cortical and sub-cortical contributions to instruction-based learning: Regional specialisation and global network dynamics. NeuroImage, 92, pp.88–100.

Hampshire, A., Hellyer, P.J., Parkin, B., Hiebert, N., MacDonald, P., Owen, A.M., Leech, R. and Rowe, J., (2016). Network mechanisms of intentional learning. Neuroimage, 127, pp.123–134.

Hampshire, A., Highfield, R.R., Parkin, B.L. and Owen, A.M., (2012). Fractionating human intelligence. Neuron, 76(6), pp.1225–1237.

Hampshire, A. and Owen, A.M., (2006). Fractionating attentional control using event-related fMRI. Cerebral Cortex, 16(12), pp.1679–1689.

Hampshire, A. and Sharp, D.J., (2015). Contrasting network and modular perspectives on inhibitory control. Trends in cognitive sciences, 19(8), pp.445–452.

Hampshire, A., Thompson, R., Duncan, J. and Owen, A.M., (2008). The target selective neural response-similarity, ambiguity, and learning effects. PLoS One, 3(6), p.e2520.

Henson, R., (2005). What can functional neuroimaging tell the experimental psychologist?. The Quarterly Journal of Experimental Psychology Section A, 58(2), pp.193–233.

Hoshi, E., Shima, K. and Tanji, J., (1998). Task-dependent selectivity of movement-related neuronal activity in the primate prefrontal cortex. Journal of neurophysiology, 80(6), pp.3392–3397.

Hsu, N.S., Jaeggi, S.M. and Novick, J.M., (2017). A common neural hub resolves syntactic and non-syntactic conflict through cooperation with task-specific networks. Brain and language, 166, pp.63–77.

Jenkinson, M., Bannister, P., Brady, J. M. and Smith, S. M. (2002) Improved Optimisation for the Robust and Accurate Linear Registration and Motion Correction of Brain Images. NeuroImage, 17(2), 825–841.

Jenkinson, M., Beckmann, C.F., Behrens, T.E., Woolrich, M.W., Smith, S.M. FSL., (2012) NeuroImage, 62:782–90.

Kan, I.P., Teubner-Rhodes, S., Drummey, A.B., Nutile, L., Krupa, L. and Novick, J.M., (2013). To adapt or not to adapt: The question of domain-general cognitive control. Cognition, 129(3), pp.637–651.

Kiesel, A., Kunde, W. and Hoffmann, J., (2006). Evidence for task-specific resolution of response conflict. Psychonomic Bulletin& Review, 13(5), pp.800–806.

Kim, C., Chung, C. and Kim, J., (2010). Multiple cognitive control mechanisms associated with the nature of conflict. Neuroscience letters, 476(3), pp.156–160.

Kim, C., Chung, C. and Kim, J., (2012). Conflict adjustment through domain-specific multiple cognitive control mechanisms. Brain research, 1444, pp.55–64.

Kunde, W. and Wühr, P., (2006). Sequential modulations of correspondence effects across spatial dimensions and tasks. Memory& Cognition, 34(2), pp.356–367.

Li, C.S.R., Huang, C., Constable, R.T. and Sinha, R., (2006). Imaging response inhibition in a stop-signal task: neural correlates independent of signal monitoring and post-response processing. Journal of Neuroscience, 26(1), pp.186–192.

Li, S., Ostwald, D., Giese, M. and Kourtzi, Z., (2007). Flexible coding for categorical decisions in the human brain. Journal of Neuroscience, 27(45), pp.12321–12330.

Li, Q., Yang, G., Li, Z., Qi, Y., Cole, M.W. and Liu, X., (2017). Conflict detection and resolution rely on a combination of common and distinct cognitive control networks. Neuroscience& Biobehavioral Reviews, 83, pp.123–131.

Linden, D.E., Prvulovic, D., Formisano, E., Völlinger, M., Zanella, F.E., Goebel, R. and Dierks, T., (1999). The functional neuroanatomy of target detection: an fMRI study of visual and auditory oddball tasks. Cerebral cortex, 9(8), pp.815–823.

Lorenz, R., Violante, I.R., Monti, R.P., Montana, G., Hampshire, A. and Leech, R., (2018). Dissociating frontoparietal brain networks with neuroadaptive Bayesian optimization. Nature communications, 9(1), pp.1–14.

Melcher, T., Falkai, P. and Gruber, O., (2008). Functional brain abnormalities in psychiatric disorders: neural mechanisms to detect and resolve cognitive conflict and interference. Brain Research Reviews, 59(1), pp.96–124.

Nieuwenhuis, S., Yeung, N., Van Den Wildenberg, W. and Ridderinkhof, K.R., (2003). Electrophysiological correlates of anterior cingulate function in a go/no-go task: effects of response conflict and trial type frequency. Cognitive, affective,& behavioral neuroscience, 3(1), pp.17–26.

Notebaert, W. and Verguts, T., (2008). Cognitive control acts locally. Cognition, 106(2),pp.1071–1080.

Nyberg, L., Marklund, P., Persson, J., Cabeza, R., Forkstam, C., Petersson, K.M. and Ingvar, M., (2003). Common prefrontal activations during working memory, episodic memory, and semantic memory. Neuropsychologia, 41(3), pp.371–377.

Rao, S.C., Rainer, G. and Miller, E.K., 1997. Integration of what and where in the primate prefrontal cortex. Science, 276(5313), pp.821–824.

Rougier, N.P., Noelle, D.C., Braver, T.S., Cohen, J.D. and O’Reilly, R.C., (2005). Prefrontal cortex and flexible cognitive control: Rules without symbols. Proceedings of the National Academy of Sciences, 102(20), pp.7338–7343.

Ruge, H. and Wolfensteller, U., (2013). Functional integration processes underlying the instruction-based learning of novel goal-directed behaviors. NeuroImage, 68, pp.162–172.

Shashidhara, S., Mitchell, D.J., Erez, Y. and Duncan, J., (2019). Progressive recruitment of the frontoparietal multiple-demand system with increased task complexity, time pressure, and reward. Journal of cognitive neuroscience, 31(11), pp.1617–1630.

Sigala, N., Kusunoki, M., Nimmo-Smith, I., Gaffan, D. and Duncan, J., (2008). Hierarchical coding for sequential task events in the monkey prefrontal cortex. Proceedings of the National Academy of Sciences, 105(33), pp.11969–11974.

Smith, S.M., (2002). Fast robust automated brain extraction. Human brain mapping, 17(3), pp.143–155.

Soreq, E., Leech, R. and Hampshire, A., (2019). Dynamic network coding of working-memory domains and working-memory processes. Nature communications, 10(1), pp.1–14.

Soreq, E., Violante, I., Hampshire, A. (2020). An Imaging Based Network Sampling Theory of Intelligence. To be published in Nature Communications.

Stiers, P., Mennes, M. and Sunaert, S., (2010). Distributed task coding throughout the multiple demand network of the human frontal–insular cortex. Neuroimage, 52(1), pp.252–262.

Stokes, M.G., Buschman, T.J. and Miller, E.K., (2017). Cognitive control. The Wiley handbook of cognitive control, p.221.

Verbruggen, F., Liefooghe, B., Notebaert, W. and Vandierendonck, A., (2005). Effects of stimulus–stimulus compatibility and stimulus–response compatibility on response inhibition. Acta psychologica, 120(3), pp.307–326

Woolgar, A., Hampshire, A., Thompson, R. and Duncan, J., (2011a). Adaptive coding of task-relevant information in human frontoparietal cortex. Journal of Neuroscience, 31(41), pp.14592–14599.

Woolgar, A., Thompson, R., Bor, D. and Duncan, J., (2011b). Multi-voxel coding of stimuli, rules, and responses in human frontoparietal cortex. Neuroimage, 56(2), pp.744–752.

Wu, T., Chen, C., Spagna, A., Wu, X., Mackie, M.A., Russell-Giller, S., Xu, P., Luo, Y.J., Liu, X., Hof, P.R. and Fan, J., (2020). The functional anatomy of cognitive control: A domain-general brain network for uncertainty processing. Journal of comparative neurology, 528(8), pp.1265–1292.

